# Benchmarking Peak Calling Methods for CUT&RUN

**DOI:** 10.1101/2024.11.13.622880

**Authors:** Amin Nooranikhojasteh, Ghazaleh Tavallaee, Elias Orouji

## Abstract

Cleavage Under Targets and Release Using Nuclease (CUT&RUN) has rapidly gained prominence as an effective approach for mapping protein-DNA interactions, especially histone modifications, offering substantial improvements over conventional chromatin immunoprecipitation sequencing (ChIP-seq). However, the effectiveness of this technique is contingent upon accurate peak identification, necessitating the use of optimal peak calling methods tailored to the unique characteristics of CUT&RUN data. Here, we benchmark four prominent peak calling tools, MACS2, SEACR, GoPeaks, and LanceOtron, evaluating their performance in identifying peaks from CUT&RUN datasets. Our analysis utilizes in-house data of three histone marks (H3K4me3, H3K27ac, and H3K27me3) from mouse brain tissue, as well as samples from the 4D Nucleome database. We systematically assess these tools based on parameters such as the number of peaks called, peak length distribution, signal enrichment, and reproducibility across biological replicates. Our findings reveal substantial variability in peak calling efficacy, with each method demonstrating distinct strengths in sensitivity, precision, and applicability depending on the histone mark in question. These insights provide a comprehensive evaluation that will assist in selecting the most suitable peak caller for high-confidence identification of regions of interest in CUT&RUN experiments, ultimately enhancing the study of chromatin dynamics and transcriptional regulation.

## Introduction

In recent years, CUT&RUN (Cleavage Under Targets and Release Using Nuclease) has emerged as a powerful alternative to traditional ChIP-seq for profiling protein-DNA interactions, particularly histone modifications. (1) This technique offers a higher signal-to-noise ratio, requires less input material, and provides a more precise mapping of protein-DNA binding sites. (2) However, as with any genomic technique, the accuracy and efficiency of data analysis heavily depend on the tools used to call peaks from the raw sequencing data. Proper peak calling is crucial for identifying regions of interest, such as transcription factor binding sites or histone modifications, but choosing the best peak calling method for a given dataset remains a challenge.

The diversity of peak calling algorithms and their underlying assumptions can lead to significantly different results when applied to the same dataset. For instance, traditional ChIP-seq peak callers like MACS2 (3) are widely used but may not be fully optimized for CUT&RUN’s unique signal characteristics. Conversely, newer tools such as SEACR (4), which was specifically designed for CUT&RUN data, claim to better exploit the sparse but strong signal profile typical of this technique. Other tools, like GoPeaks (5) and LanceOtron (6), bring their own strengths, such as robust binomial modeling or machine learning-based predictions, further complicating the decision of which tool is best suited for analyzing CUT&RUN datasets.

The aim of this study is to benchmark four peak calling methods, MACS2, SEACR, GoPeaks, and LanceOtron on their ability to accurately and efficiently detect peaks from CUT&RUN data. We focus on CUT&RUN experiments profiling three histone marks, H3K4me3, H3K27ac, and H3K27me3, using two replicates per mark from mouse brain tissue that are generated in-house along with four samples with different number of replicates from the 4D Nucleome database (a total of 15 samples across the three histone marks). By comparing the number of peaks called, peak length, signal enrichment, and reproducibility across the methods, we aim to determine which tool is the most effective for CUT&RUN data analysis. Our findings will provide insights on parameters to consider while selecting peak calling tools and help improve the reliability of CUT&RUN experiments in studying chromatin dynamics and transcriptional regulation.

In this paper^1^, we address the following key questions: (1) Which peak calling method identifies the highest number of biologically relevant peaks? (2) How consistent are these peaks across replicates? (3) What are the trade-offs between peak sensitivity and specificity across different tools? By systematically comparing these tools on the same dataset, we seek to provide insights into which peak calling method is best suited for high-confidence peak identification in CUT&RUN experiments.

## Materials and Methods

### Sample Collection and CUT&RUN Experimental Design

#### Mouse Samples

Adult C57BL6 mice were used for this study. Fresh brain tissue was obtained from female mice of 8-10 weeks of age. All animal samples obtained in this study complied the relevant ethical regulations approved by the institutional ethics comimittee and Research Ethics Board at the University Health Network (UHN). The chromatin profiling in this study targeted three specific histone modifications: H3K4me3; associated with active transcription start sites and promoters, H3K27ac; a marker for active enhancers and promoters and H3K27me3; related to repressive chromatin domains and silenced genes. Two biological replicates were prepared for each histone mark. These histone marks were assayed using anti-H3K4me3 (Abcam, Cat. ab8580), anti-H3K27ac (Abcam, Cat. ab4729) and anti-H3K27me3 (Diagenode, Cat. C15410069). The most commonly used and optimized CUT&RUN protocol (1) was used for all samples, ensuring minimal background noise and high signal-to-noise ratio typical of CUT&RUN experiments.

#### 4D Nucleome Datasets

For this study, we obtained publicly available CUT&RUN datasets from the 4D Nucleome Data Portal (https://data.4dnucleome.org/). We selected data for four specific samples, each with technical replicates, for which data was available for all three histone marks investigated in this study: H3K4me3, H3K27ac, and H3K27me3. The files were retrieved by accessing the unique dataset identifiers listed in the 4D Nucleome database. Each dataset was then systematically downloaded using the portal’s API, ensuring data integrity through verification of checksum values. The BAM files were subsequently processed for quality control and peak calling to facilitate further analysis and integration in our research. All data processing was conducted on UHN’s high-performance computing infrastructure, ensuring reproducibility and consistency across the analyses.

**Table 1** shows details of the samples used in this study.

**Table 1.** Sample IDs, number of replicates, origin, species and source of data.

| Sample Name | Replicate | Origin | Species | Source |
| --- | --- | --- | --- | --- |
| GM12878_R1 | R1 | Lymphoblast | Human | 4DNucleome |
| GM12878_R2 | R2 | Lymphoblast | Human | 4DNucleome |
| H1_ESC_R1 | R1 | Embryonic Stem Cells (ESC) | Human | 4DNucleome |
| H1_ESC_R2 | R2 | Embryonic Stem Cells (ESC) | Human | 4DNucleome |
| H1_ESC_R3 | R3 | Embryonic Stem Cells (ESC) | Human | 4DNucleome |
| H1_ESC_R4 | R4 | Embryonic Stem Cells (ESC) | Human | 4DNucleome |
| H1_ESC_R5 | R5 | Embryonic Stem Cells (ESC) | Human | 4DNucleome |
| HFFc6_R1 | R1 | Foreskin Fibroblast (HFFc6) | Human | 4DNucleome |
| HFFc6_R2 | R2 | Foreskin Fibroblast (HFFc6) | Human | 4DNucleome |
| HFFc6_R3 | R3 | Foreskin Fibroblast (HFFc6) | Human | 4DNucleome |
| HFFc6_R4 | R4 | Foreskin Fibroblast (HFFc6) | Human | 4DNucleome |
| K562_R1 | R1 | Myelogenous Leukemia | Human | 4DNucleome |
| K562_R2 | R2 | Myelogenous Leukemia | Human | 4DNucleome |
| Mouse_Brain_R1 | R1 | Brain | Mouse | In-house* |
| Mouse_Brain_R2 | R2 | Brain | Mouse | In-house* |

\* Mouse brain samples were generated in-house (Princess Margaret Cancer Centre) and data for the other samples are obtained from the 4DNucleome portal.

### CUT&RUN Protocol

The CUT&RUN procedure was performed following the standard protocol described by Meers et al. (1). Fresh brain tissue was homogenized, and nuclei were isolated using a lysis buffer optimized for brain tissue, followed by incubation with Concanavalin A-coated magnetic beads. The isolated nuclei were then incubated overnight at 4°C with the respective antibodies targeting the histone marks H3K4me3, H3K27ac, H3K27me3 and IgG. Antibody concentrations were optimized according to the manufacturer’s recommendations. Following antibody binding, the nuclei were treated with Protein A/G-MNase fusion protein to cleave the chromatin at the antibody-bound sites. After MNase activation, the DNA fragments were released and collected for downstream sequencing.

### Sequencing and Data Processing

#### Library Preparation and Sequencing

CUT&RUN libraries were prepared from the released DNA fragments using the NEBNext Ultra II DNA Library Prep Kit (New England BioLabs). The libraries were sequenced using Illumina NovaSeq 6000 with paired-end 50 base pair (50-PE) reads at a sequencing depth of approximately 40 million reads per sample.

#### Data Preprocessing

Raw sequencing reads were processed using the nf-core/cutandrun pipeline (v3.2.2) with default settings (1). FASTQ files were quality checked using FastQC (v0.11.9), followed by adapter trimming using Trim Galore (v0.6.10). Reads were then aligned to either the mm10 mouse or hg38 human reference genome using Bowtie2 (v2.4.1) with default parameters, including the use of both target and spike-in genomes. The resulting BAM files were sorted and indexed with SAMtools (v1.11), and duplicate reads were marked using Picard (v3.1.1).

### Peak Calling Methods

Four peak calling methods were benchmarked in this study: MACS2, SEACR, GoPeaks, and LanceOtron. Each method was run using default parameters unless otherwise specified, with the goal of determining which method most efficiently detects peaks in CUT&RUN datasets. This study included both mouse (mm10) and human (hg38) samples. Below, we describe the peak calling parameters used for each tool.

MACS2 (Model-based Analysis of ChIP-Seq) (MACS2 v2.2.9.1) was used for peak calling on BAM files aligned to both the mm10 and hg38 genomes. For each BAM file, peaks were called with the following parameters: paired-end BAM format (-f BAMPE), genome sizes set to 2.7e9 (mm10) and 3.1e9 (hg38), and a false discovery rate cutoff of 0.05 (-q 0.05). For H3K27me3-marked samples, the –broad option was applied to identify broad peaks, while no additional broad peak option was specified for other histone marks.

SEACR (Sparse Enrichment Analysis for CUT&RUN) (SEACR v1.3) was used to call peaks by first converting BAM files to bedGraph format with BEDTools genomecov. (7) SEACR was then run in stringent mode for both mm10 and hg38 alignments, with a false discovery rate threshold of 0.01, generating output files in the stringent peak format for each genome.

GoPeaks (GoPeaks v1.1) peak calling was performed on BAM files aligned to mm10 and hg38. BAM files were processed in batches, excluding index (.bai) files. For H3K27me3-marked samples, the –mdist 3000 and –broad options were applied to account for the broad distribution of this histone mark, while other histone marks used –mdist 1000. Peak files in BED format were generated and saved for each genome alignment, with an FDR threshold of 0.01.

LanceOtron (LanceOtron v1.2.7) used bigWig format as input. For each BAM file aligned to either mm10 or hg38, a bigWig file was generated using bamCoverage from Deeptools package (8), which was then used as input for LanceOtron to identify peaks. Peaks were called and saved in BED format, with bigWig files moved to a separate directory and BED files stored in the output directory.

BigWig files were processed using bamCoverage, and peaks were loaded into R as GRanges objects (9). Peak files in either BED, narrowPeak, or broadPeak formats were imported for each tool. Signal extraction from the bigWig files was performed over the peak regions, and background noise was estimated using randomly selected regions without peaks.

The SNR was calculated by dividing the signal values by the mean background noise for each condition, and the results were saved for further analysis.

### Signal-to-Noise Ratio (SNR) Calculation

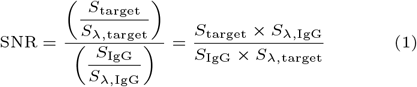

- *S*_target_: Number of reads mapped to the genome (or target regions) in the target antibody sample.
- *S*_*λ*,target_: Number of reads mapped to the spike-in DNA (lambda DNA) in the target antibody sample.
- *S*_IgG_: Number of reads mapped to the genome (or target regions) in the IgG control sample.
- *S*_*λ*,IgG_: Number of reads mapped to the spike-in DNA (lambda DNA) in the IgG control sample.

### Generation of Consensus Peak Dataset as the Ground Truth

To evaluate the performance of each peak calling method in identifying true peaks, a consensus peak set was constructed for each sample. The consensus peak set represents a rigorous and highly confident subset of peaks, comprising only those peaks that are consistently identified by a minimum of three peak calling methods evaluated in this study. By requiring a peak to be called at least by three of the four distinct methods, the resulting consensus set is derived using an exceptionally stringent criterion, enhancing the reliability of the identified peaks. Such a consensus approach, albeit conservative, minimizes false positives, thereby yielding a high-confidence peak dataset that can serve as a benchmark for subsequent evaluations.

We first identified peaks from all possible combinations of three peak callers, generating four sets of consensus peaks. The union of these overlapping regions was defined as the final consensus peak set. We then evaluated each individual peak caller’s performance against this consensus using three metrics: precision (proportion of called peaks that overlap with consensus peaks), recall (proportion of consensus peaks detected by each caller), and F1 score (harmonic mean of precision and recall). The analysis was visualized using Venn diagrams to display the overlap between different peak callers and the summary of these metrics were plotted for each peak calling method across three different marks separately.

### Precision, Recall Compositional Normalization and Ternary Plot Visualization

The precision and recall scores derived from each peak-calling method were normalized to represent the relative contributions of three histone modifications (H3K27ac, H3K4me3, and H3K27me3) in a ternary plot. Specifically, let *p*_*i,j*_ denote the precision score for histone mark *i* in replicate *j*, where *i* ∈ {H3K27ac, H3K4me3, H3K27me3}, and *j* represents the individual replicates for each condition. The normalized score *q*_*i,j*_ for each histone mark *i* was computed as follows:

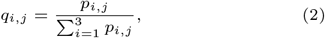

where the denominator represents the sum of precision scores across the three histone marks for each replicate, ensuring that the normalized scores for each replicate sum to unity:

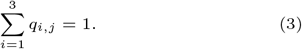

This compositional normalization method allows for visualization of the relative precision of the peak-calling methods across the three histone marks, highlighting their differential contributions. The resulting ternary plot provides a comprehensive overview of the performance of the peak-calling methods, enabling comparison across multiple conditions and replicates.

## Results

### Peak Comparison and Evaluation

#### Metrics for Benchmarking

The performance of each peak caller was evaluated based on the following criteria:

1. **Total Number of Peaks**: The total number of peaks called by each method was quantified to assess the sensitivity of each peak caller in identifying potential regions of interest across the dataset.
2. **Peak Length Distribution**: The distribution of peak lengths was analyzed for each method to compare peak length patterns and identify differences in resolution and detection of broad versus narrow peaks across the genome.
3. **Signal-to-Noise Ratio**: Enrichment of signal over background noise was evaluated by analyzing the read density across peaks. This metric helps in determining each method’s ability to distinguish true peaks from background noise, thus indicating overall signal clarity.
4. **Precision, Recall, and F1 Score Metrics**: These metrics were calculated by comparing identified peaks against a reference set of high-confidence peaks. Precision measures the fraction of true positives among all called peaks, recall measures the fraction of true positives among actual peaks, and the F1 Score provides a balance between precision and recall.
5. **Overlap Analysis Between Methods**: Pairwise comparisons were conducted to assess the overlap in peak calls across different methods. This analysis provided insights into the consistency and specificity of each peak caller.
6. **Overlap Analysis Between Replicates**: The reproducibility of peaks between biological replicates was assessed for each method using the Irreproducible Discovery Rate (IDR). IDR analysis determined the consistency of peak detection between replicates, which is crucial for assessing method reliability.

#### Total Number of Peaks Called

The total number of peaks called by each peak caller, MACS2, SEACR, GoPeaks, and LanceOtron showed marked differences across the histone marks H3K4me3, H3K27ac, and H3K27me3. LanceOtron identified the highest number of peaks across all histone marks, with an average of 110,876 for H3K4me3, 367,023 for H3K27ac, and 281,545 for H3K27me3. This was consistent across both biological replicates for each histone mark, indicating LanceOtron’s sensitivity to the sparse, high-intensity signal typical of CUT&RUN datasets.

In contrast, GoPeaks called the fewest peaks, with significantly lower numbers across all histone marks: 24,623 for H3K4me3, 23,280 for H3K27ac, and 21,684 for H3K27me3. The peak counts from MACS2 and SEACR were intermediate, with MACS2 detecting 34,063 peaks on average for H3K4me3, 39,873 for H3K27ac, and 28,861 for H3K27me3, while SEACR identified 31,976, 182,261, and 583,414 peaks for the same marks, respectively. These trends suggest that GoPeaks is the most stringent in peak detection, potentially at the expense of missing biologically relevant but weaker peaks, whereas LanceOtron captures a broader spectrum of peaks, which may include more false positives. **(Figure 1a and 2a)**

**Fig. 1.**
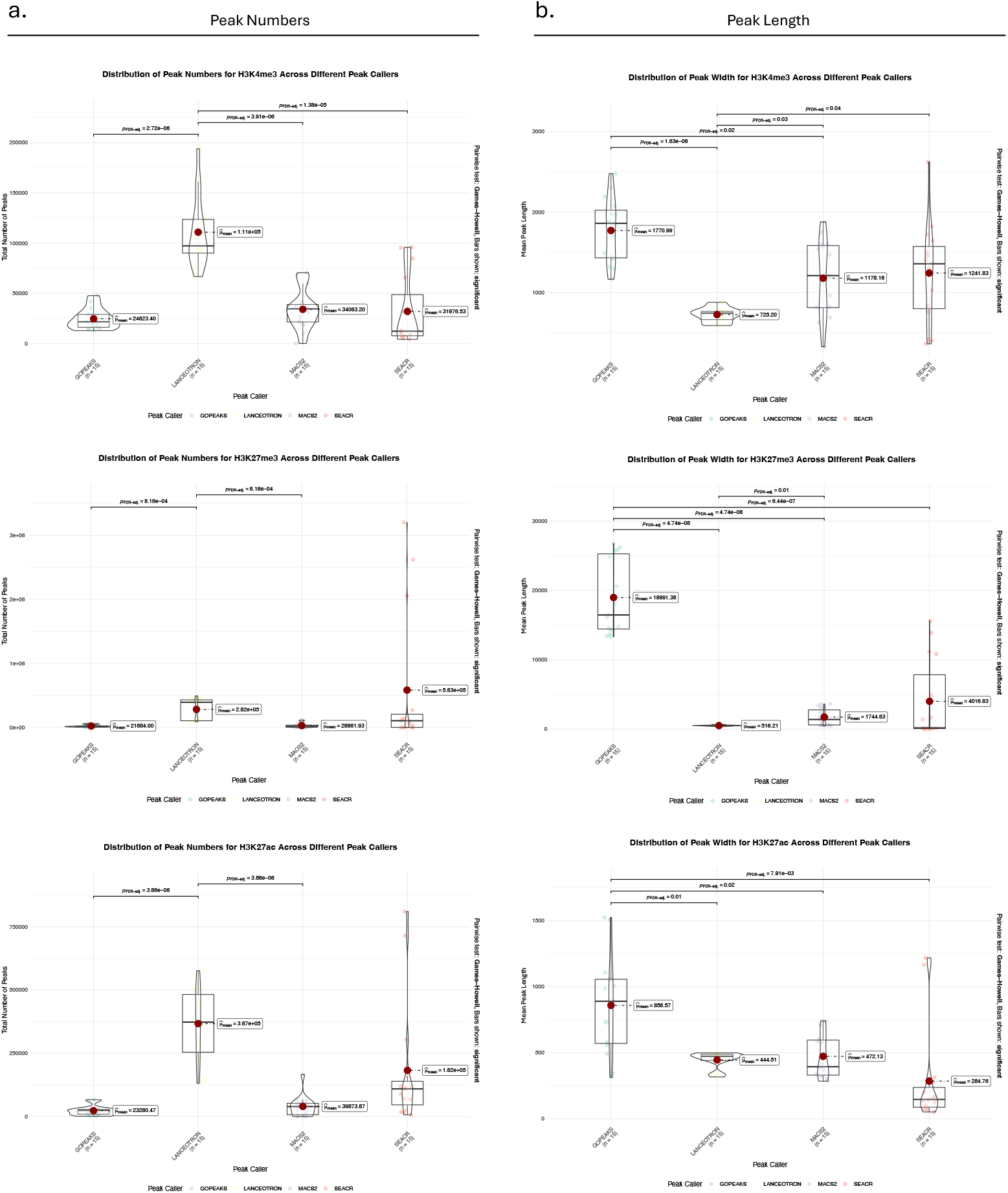
Distribution of peak numbers and length in three histone marks using four peak calling methods.

**Fig. 2.**
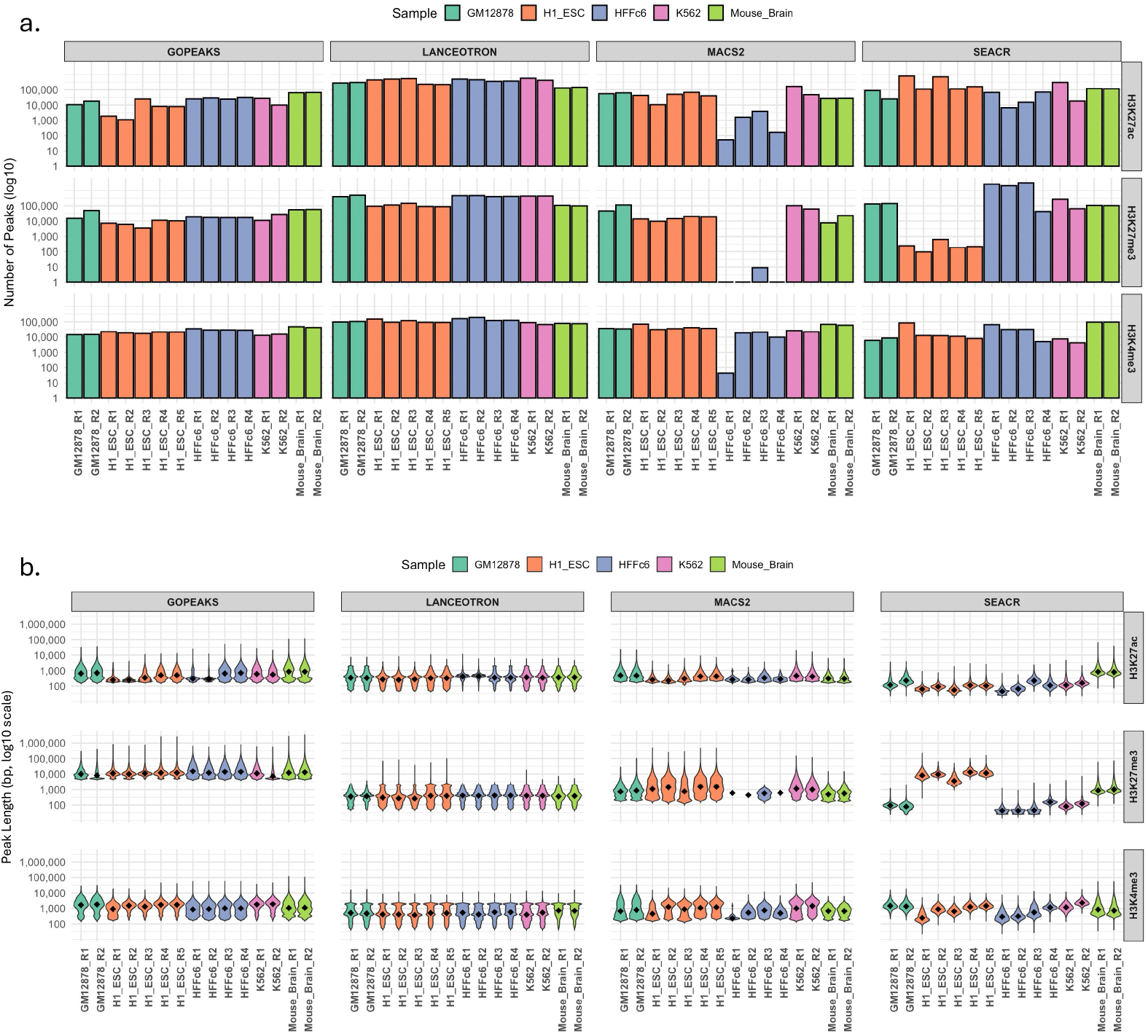
Distribution of peak numbers and length across all individual samples with their replicates using all the four peak calling methods.

### Peak Length Distribution

The distribution of peak lengths varied significantly between the peak callers. GoPeaks produced consistently longer peaks compared to MACS2, SEACR, and LanceOtron across all histone marks. The mean peak length for H3K4me3 using GoPeaks was 1,770 bp, while LanceOtron produced much narrower peaks with a mean length of 725 bp. For H3K27ac, GoPeaks’s peak lengths were similarly broader, with a mean length of 858 bp compared to the narrowest peaks in SEACR with the mean of 284 bp. This trend continued with H3K27me3, where GoPeaks identified much broader repressive regions, averaging 18,991 bp in length, while LanceOtron called the narrowest peaks at 516 bp. With the exception of the H3K27ac mark, MACS2 and SEACR showed peak length distributions that were intermediate between GoPeaks and LanceOtron, with MACS2 detecting mean peak lengths of 1,178 bp for H3K4me3, 472 bp for H3K27ac, and 1,744 bp for H3K27me3, while SEACR detected 1,241 bp, 284 bp, and 4,016 bp for the respective marks. **(Figure 1b and 2b)**

### Signal-to-Noise Ratio (SNR)

To assess the signal enrichment of each peak caller, we examined the signal-to-noise ratio (SNR) by plotting read density over the identified peak regions. SNR analysis was performed using bigWig files and peak data from all the peak calling tools for the histone marks H3K27ac, H3K27me3, and H3K4me3 in all replicates. **(Figure 3)**

**Fig. 3.**
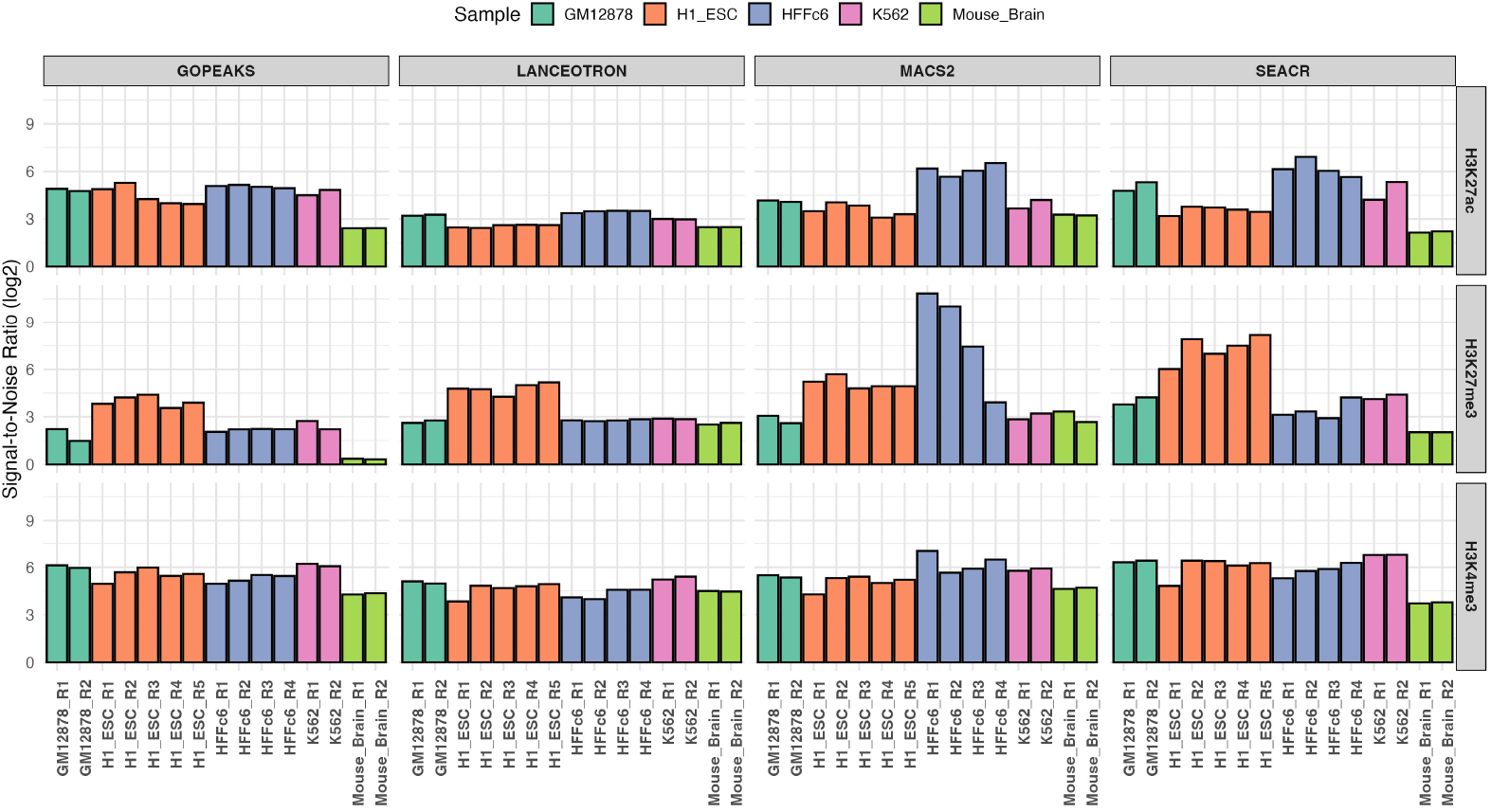
Signal to noise ratio (SNR) across all marks using four peak calling methods.

SNR analysis revealed distinct differences in sensitivity for detecting peaks across three histone marks (H3K27ac, H3K27me3, and H3K4me3). SEACR consistently achieved the highest SNR across all marks, underscoring its superior sensitivity and suitability for high-confidence peak detection, particularly in complex chromatin contexts involving both active and repressive marks.

MACS2 demonstrated high SNR for active histone marks (H3K27ac and H3K4me3), indicating strong signal enrichment for transcriptionally active regions. GoPeaks and LanceOtron provided moderate, stable SNR values across all marks, reflecting consistent performance but with lower sensitivity than SEACR, especially for marks with subtle signal patterns like H3K27me3.

### Overlap Analysis Between Methods

Figure 4. presents Venn diagrams illustrating the overlap among the four peak calling methods for each sample. These diagrams, generated separately for each histone mark (H3K27ac, H3K4me3, and H3K27me3) and each sample (H1 ESC, K562, HFFc6, GM12878, and Mouse-Brain), provide key insights into the similarities and discrepancies in peak detection across the methods.

The Venn diagrams for H3K4me3 in the Mouse Brain and H1 ESC samples show a considerable overlap among all four peak calling methods, with 43,378 (37%) and 23,502 (12%) peaks, respectively, shared among the four methods.

**Fig. 4.**
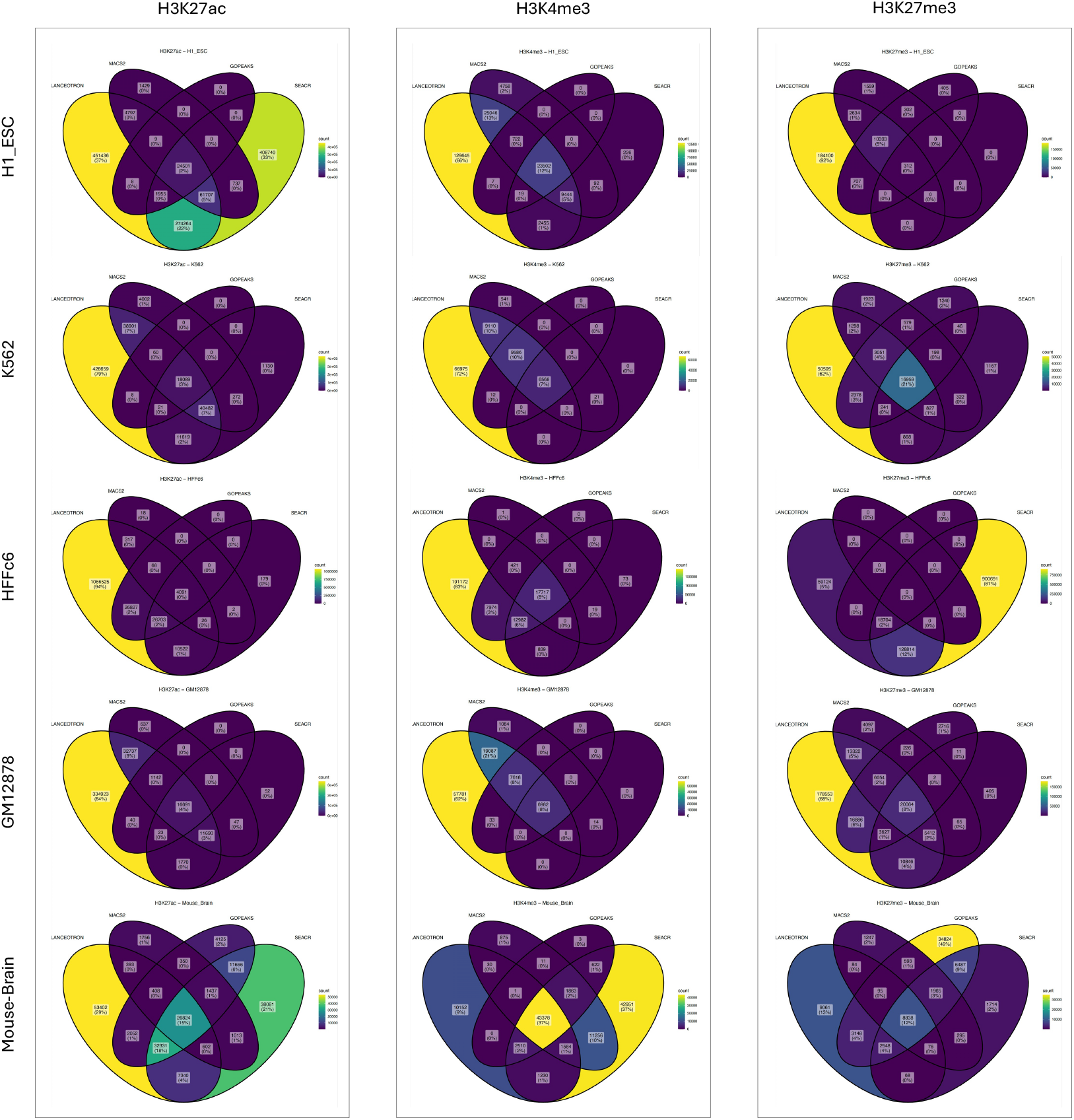
Overlap of peaks called using all the four peak calling methods across the samples.

For H3K27me3, the highest percentage of overlapping peaks was observed in K562 with 16,959 (21%), followed by 8,838 (12%) in Mouse Brain and 20,064 (8%) in GM12878.

For H3K27ac, the percentage of shared peaks among all four methods did not exceed 4%, with the exception of the Mouse Brain sample, which had 26,824 (15%) shared peaks.

A consistent pattern across histone marks and samples was observed, with GoPeaks detecting fewer peaks. Conversely, LanceOtron tended to call the highest number of peaks, generating the highest number of peaks in 12 out of 15 samples across all three histone marks.

The overlap analysis reveals that, despite the inherent differences among the peak callers, a substantial number of peaks are shared across 2-3 methods, particularly for the active histone marks H3K27ac and H3K4me3. The overlap for H3K27me3, a repressive mark, is notably lower, which might reflect the broader and more variable nature of these peaks. **(Figure 4 and Table 2)**

**Table 2.**
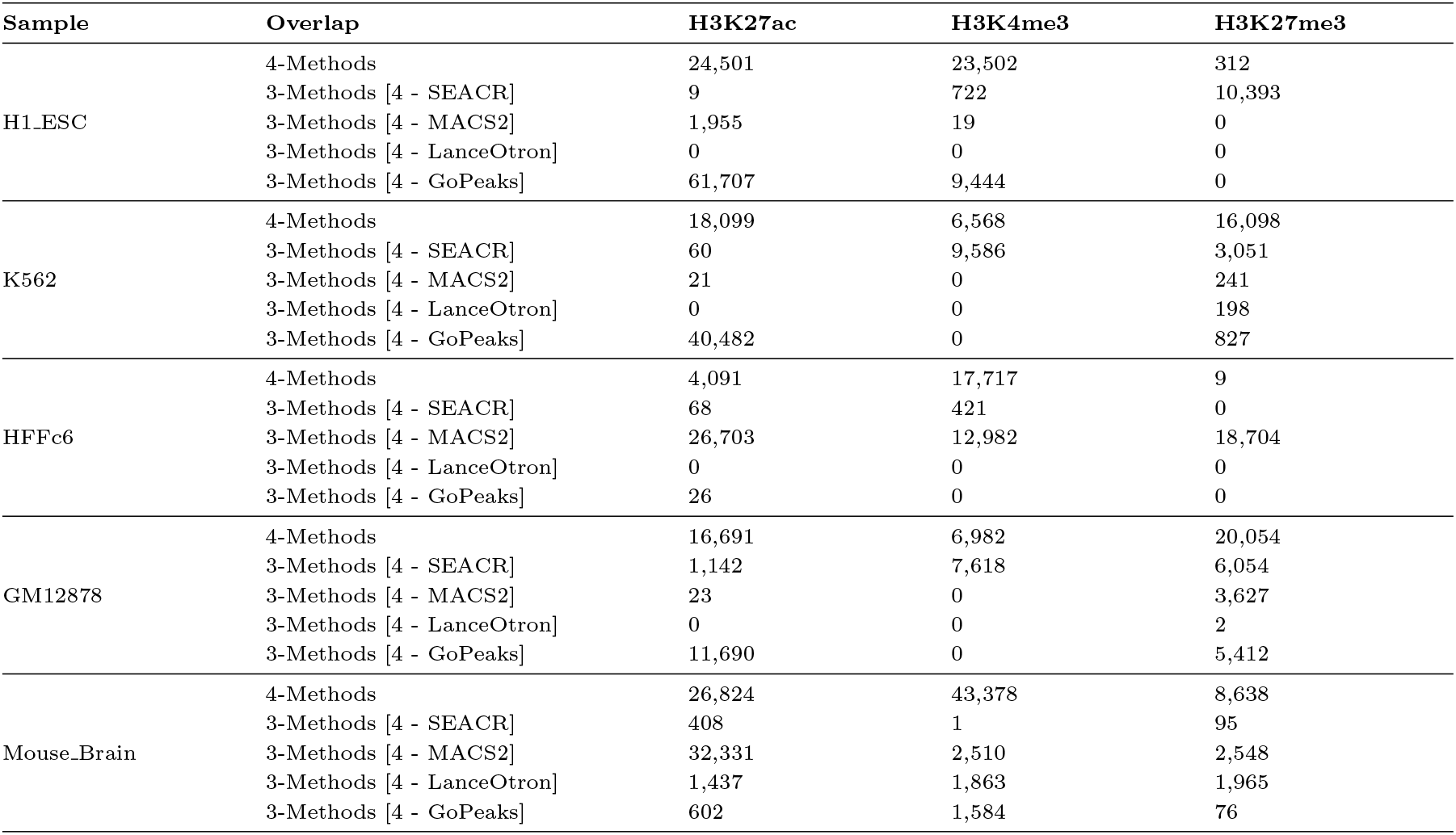
Number of overlapping peaks among four or three peak-calling methods across various samples and histone marks.

### Precision, Recall, and F1 Score Metrics

Using the consensus peak set as the reference, we assessed the precision, recall, and F1 score of each peak calling tool. These metrics were calculated by comparing identified peaks against a reference set of high-confidence peaks, derived as described in the Methods section. Precision was defined as the proportion of peaks called by each method that overlap with the consensus set, while recall was defined as the fraction of consensus peaks successfully identified by each method. The F1 score, being the harmonic mean of precision and recall, provided a balanced measure that considered both false positives and false negatives. **Figure 5** shows the performance of each peak calling method in terms of precision across three different histone marks (H3K4me3, H3K27ac, and H3K27me3). Among the methods, GoPeaks exhibited the highest precision for H3K4me3 and H3K27ac, while SEACR performed comparably for H3K27me3. Conversely, LanceOtron displayed the lowest precision across most histone marks, which may indicate its higher rate of false positives, particularly for H3K4me3 and H3K27ac.

**Fig. 5.**
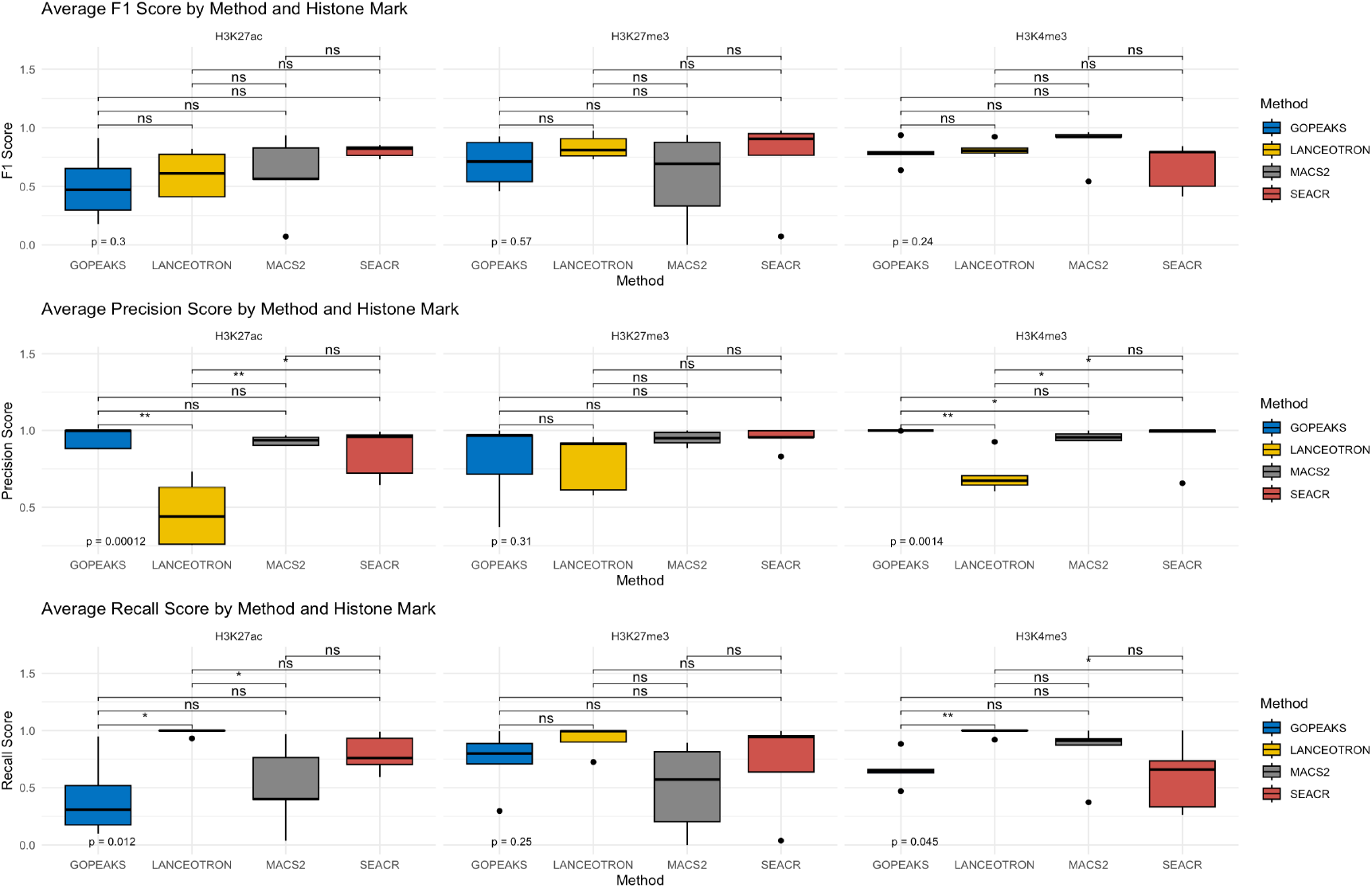
Precision, recall and F1 score for all four peak calling methods based on the histone mark tested.

In terms of recall, LanceOtron and SEACR exhibited superior performance compared to MACS2 and GoPeaks for all three histone marks, with LanceOtron achieving the highest recall values for H3K27me3. This trend was consistent for LanceOtron suggesting the method to be more sensitive in detecting true positives, albeit at the potential expense of a higher number of false positives, as suggested by its relatively lower precision values.

The F1 scores illustrated in **Figure 5** for each peak caller is summarizing the trade-off between precision and recall. SEACR had the highest F1 scores for H3K27ac and H3K27me3, highlighting its balanced performance in identifying true peaks with reasonable precision and recall. MACS2 performed best for H3K4me3, while GoPeaks and LanceOtron showed comparable F1 scores.

Ternary plots presented in **Figure 6** indicate a clear distinction in performance based on the histone mark being analyzed. Higher performance of GoPeaks in precision makes it a suitable choice for active marks like H3K4me3, while its higher sensitivity makes it one of the top choices for broader peaks such as those found in H3K27me3. On the other hand, MACS2 and SEACR’s balanced performance in terms of precision and recall could make them suitable for any given histone mark regardless of their peak length. These metrics provide a comprehensive understanding of each method’s ability to accurately identify regions of interest, balancing sensitivity and specificity.

### Overlap Analysis Between Replicates

In the irreproducible discovery rate (IDR) analysis of mouse brain samples, we evaluated peak reproducibility across four peak-calling methods, MACS2, GoPeaks, LanceOtron, and SEACR by comparing two replicates. MACS2 demonstrated good concordance between replicates, with a moderate spread of IDR values, indicating it captures peaks with a balanced level of reproducibility. GoPeaks displayed high reproducibility, with most significant peaks clustering tightly along the diagonal, reflecting robust detection of consistent peaks across replicates. LanceOtron, while showing general concordance, exhibited greater variability in IDR values, suggesting it may capture more variable peaks with less reproducibility than other methods. SEACR identified a large number of peaks, with a high proportion of peaks showing reproducibility; however, it exhibited a broader distribution, indicating a higher detection rate of potentially reproducible peaks but with some variability **(Figure 7)**.

**Fig. 6.**
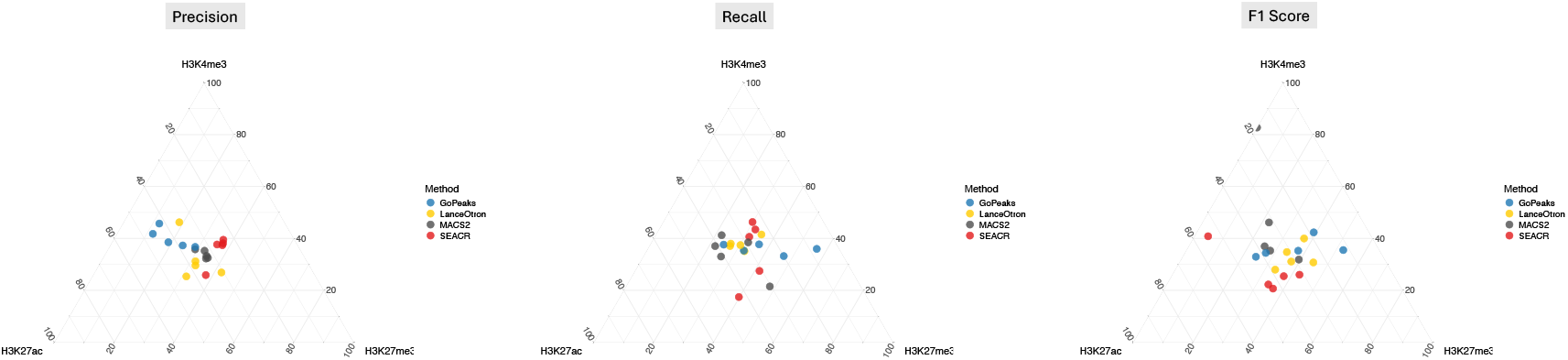
Precision, recall and F1 score for all the four peak calling methods based on the histone mark tested.

**Fig. 7.**
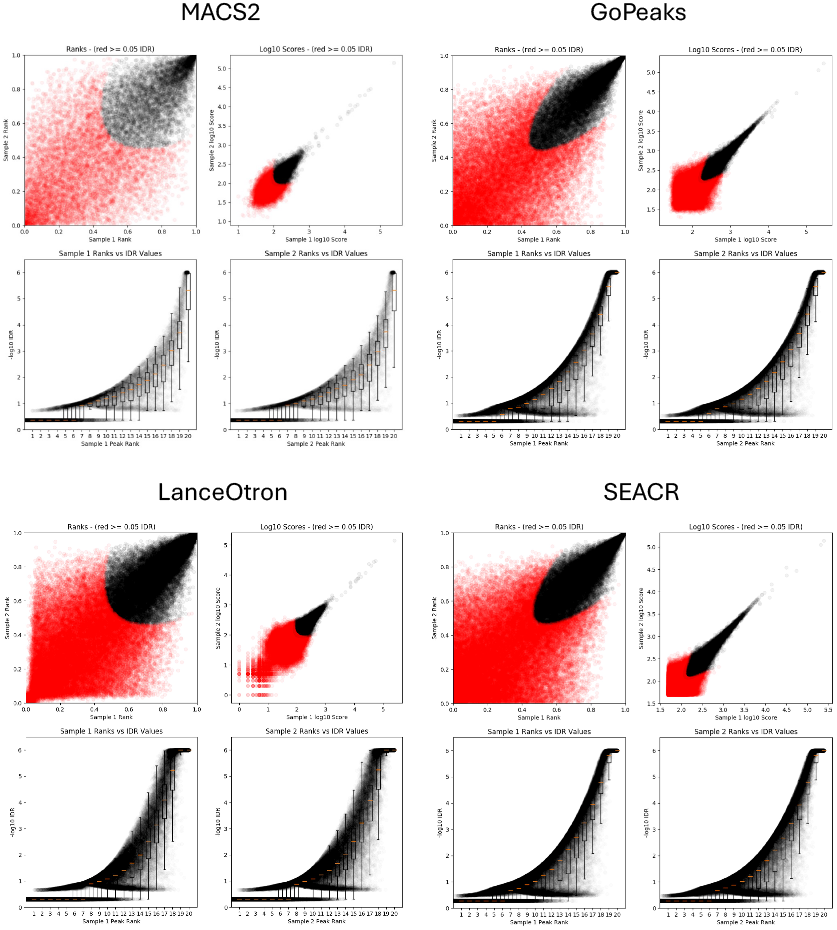
IDR analysis for the overlap between the replicates using all the four peak calling methods.

## Discussion

This study provides a comprehensive benchmarking of four peak calling methods, MACS2, SEACR, GoPeaks, and LanceOtron for the analysis of CUT&RUN datasets, focusing on the histone modifications H3K4me3, H3K27ac, and H3K27me3. Our results highlight significant variability in the performance of these tools, emphasizing the importance of selecting an appropriate peak caller for the unique characteristics of CUT&RUN data. The diversity of performance metrics, including peak number, peak length distribution, signal-to-noise ratio, and overlap across replicates, underscores the complexity of accurately identifying regions of interest and the need for tailored peak calling strategies.

The key finding of our analysis is the distinct behavior of each peak caller, reflecting differences in their sensitivity, precision, and overall efficacy. LanceOtron exhibited the highest sensitivity across all histone marks, consistently calling the largest number of peaks. However, its lower precision suggests that a substantial proportion of these peaks might represent false positives. This trade-off between sensitivity and specificity is a fundamental consideration when choosing a peak caller for a given experimental goal. For exploratory studies requiring maximal peak discovery, LanceOtron may be advantageous, while for high-confidence peak detection, additional filtering or consensus approaches may be necessary. In contrast, GoPeaks displayed a more stringent peak calling approach, identifying fewer peaks but achieving higher precision, particularly for active histone marks such as H3K4me3 and H3K27ac. This indicates that GoPeaks is more conservative in identifying true positives, making it suitable for studies prioritizing specificity over sensitivity. However, the lower recall observed for GoPeaks implies that biologically relevant peaks with weaker signals might be missed, especially for broader repressive marks like H3K27me3.

SEACR and MACS2 showed intermediate behaviors, with SEACR demonstrating balanced performance in terms of precision and recall, particularly for H3K27ac and H3K27me3. This makes SEACR an attractive option for CUT&RUN data, where both sensitivity and precision are crucial. MACS2, a tool initially developed for ChIP-seq, also performed well for H3K4me3, suggesting its applicability in CUT&RUN despite its original design for a different type of data.

Our overlap analysis between peak callers and biological replicates further emphasizes the variability inherent in peak detection across different methods. The consensus approach employed in this study, which requires a peak to be called by at least three of the four methods, provides a high-confidence set of peaks that can mitigate false positives and negatives.

The SNR analysis and peak length distribution also revealed method-specific biases. GoPeaks consistently identified broader peaks, particularly for H3K27me3, which may be advantageous when analyzing repressive chromatin domains that exhibit diffuse signal patterns. On the other hand, LanceOtron’s narrower peaks might be more suitable for defining sharp boundaries of active chromatin regions. These differences suggest that the optimal peak caller may vary depending on the specific histone mark and biological context under investigation.

In conclusion, our benchmarking analysis illustrates that there is no one-size-fits-all solution for peak calling in CUT&RUN data. Instead, the choice of peak caller should be guided by the specific experimental objectives, whether maximizing sensitivity, ensuring high precision, or achieving a balance between the two. Combining multiple peak callers or utilizing consensus-based approaches may offer a robust strategy for achieving high-confidence results, particularly in studies aiming to delineate complex chromatin landscapes. Future improvements in peak calling algorithms, potentially incorporating machine learning and integrative approaches, hold promise for enhancing the reliability and accuracy of peak detection in CUT&RUN and other next-generation sequencing assays.

## Data Availability

All the CUT&RUN datasets related to this work will be available upon publication. All the 4D Nucleome datasets can be obtained from the 4D Nucleome Data Portal (https://data.4dnucleome.org/).

## Author Contributions Statement

A.N. and E.O. conceived the experiments. G.T. conducted the experiments. A.N. and E.O. analyzed the results. A.N. and E.O. wrote and reviewed the manuscript.

## Acknowledgments

The authors thank the Princess Margaret Genomics Centre for their assistance with sample sequencing. This work was supported in part by funding from the Canadian Institutes of Health Research (CIHR — PJT-173283).

## Footnotes

1 https://github.com/OroujiLab/CUTandRun_Peak_Calling/

